# Comparison of MR coil options for concurrent TMS-fMRI

**DOI:** 10.1101/2024.05.15.594454

**Authors:** J. B. Jackson, C. L. Scrivener, M. Mada, A. Woolgar

## Abstract

Concurrent transcranial magnetic stimulation (TMS) with functional magnetic resonance imaging (fMRI) is increasingly being used to study how TMS affects neural processing in local and remote connected brain regions. However, there are many technical challenges that arise when operating this unique combination of techniques. One such challenge is MR head coil compatibility and the image resolution that can be achieved. Here we compare the temporal-signal-to noise-ratio (tSNR) between 3 different MR head coils that can be used with TMS. We show that the 7-channel TMS-dedicated surface coils result in a very high tSNR directly under the TMS and the supplementary coil in comparison to the TxRx and the Magnetica. However, there is low tSNR elsewhere. This field inhomogeneity may not be suitable for all research questions, for example where the aim is to look at distributed neural responses. In these cases, an MR coil with a more homogeneous tSNR such as the Magnetica, may be more appropriate.

## Introduction

Transcranial magnetic stimulation (TMS) combined with functional magnetic resonance imaging (fMRI) presents a unique opportunity to concurrently perturb and measure brain activity yielding powerful causal insights into brain function. Recent years has seen a resurgence of interest in this technique, including development of new TMS-dedicated MR coils. Here we compared the temporal-signal-to-noise-ratio (tSNR), a commonly used measure as a proxy for image quality (Triantafyllou et al., 2005; Liu, 2016; Colizoli, De Gee, Van Der Zwaag, & Donner, 2020), between 3 MR head coils:

1. Conventional TxRx 1-channel birdcage coil (Siemens).
2. Oversize 8-channel coil (Magnetica).
3. Two 7-channel TMS-dedicated MR surface coils (Navarro de Lara et al., 2015, sold by MagVenture).

### Data acquisition and analysis

We acquired data with and without a TMS coil positioned over a circular phantom in a Siemens 3T Prisma-fit MRI scanner using a T2*-weighted echo planar imaging acquisition sequence (TE = 30 ms, MB factor = 1, no in-plane acceleration, 18 interleaved slices, phase encoding anterior to posterior, transversal orientation, slice thickness 3 mm, voxel size 3 mm x 3 mm, 25% distance factor, flip angle 78 degrees). For each MR coil and each voxel, the tSNR was defined as the average signal over its standard deviation through time (250 volumes per run). Calculations for tSNR were performed using FSL (v5.0.9).

A MagVenture MR-compatible air-cooled figure-of-eight TMS coil, attached to a MagPro XP stimulator, was held in position using a coil holder (MagVenture) placed in the bore. In the case of the surface coil set up, the TMS coil was mounted on top of one surface coil, and the second surface coil was secured to the coil holder using an in-house custom-built flexible attachment. For both the TxRx and the Magnetica head coil setups, the TMS coil was positioned between the phantom and the MR head coils in the first run and was removed for the second run.

## Results

The TxRx and Magnetica head coils demonstrate a more homogenous tSNR than the TMS-dedicated MR surface coils, with a higher maximum and mean tSNR for the Magnetica compared to the TxRx coil (Figure 1). The TMS-dedicated surface coils demonstrate very high tSNR directly underneath the TMS coil (and supplementary coil) but lower signal elsewhere.

**Figure 1.**
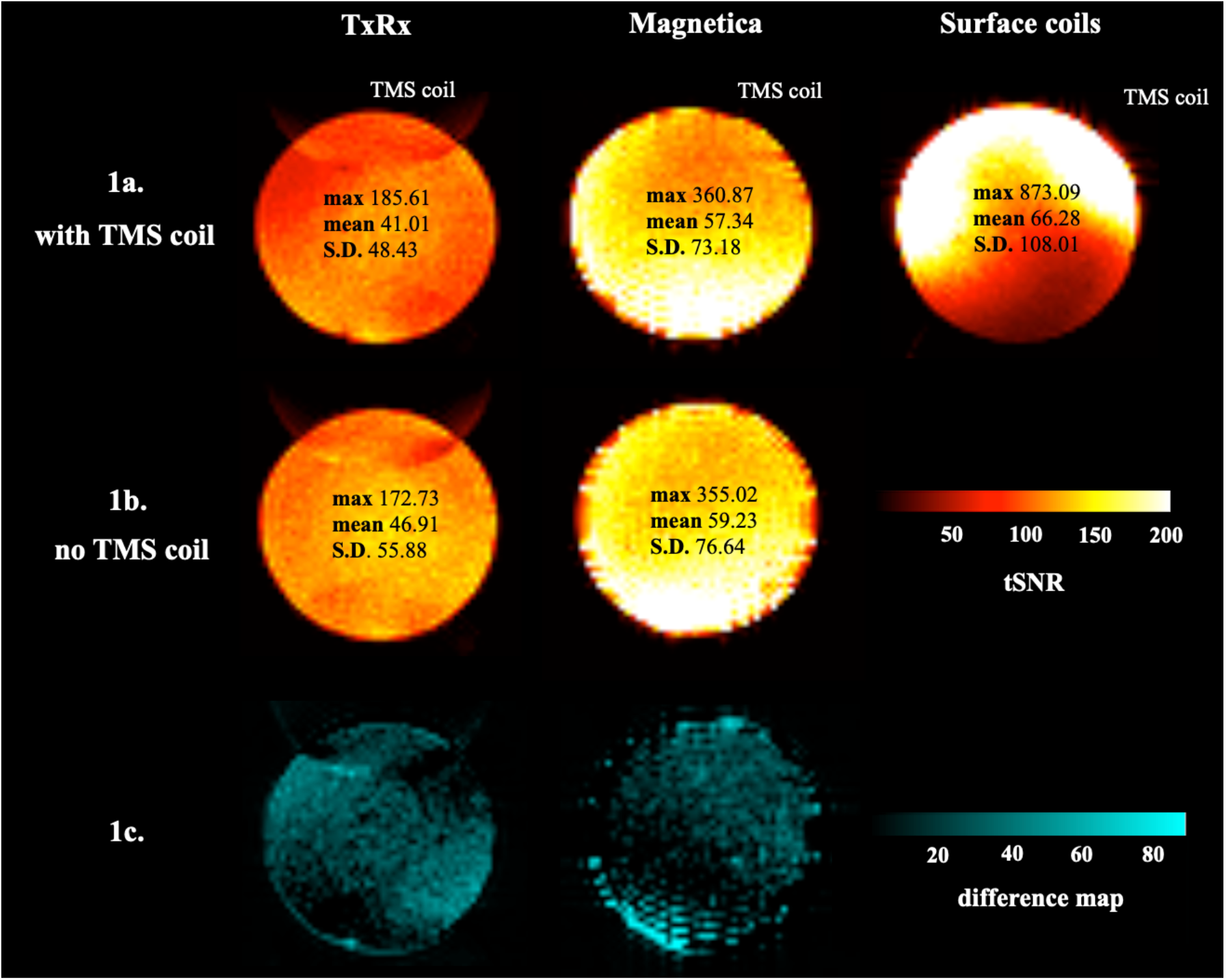
TSNR maps for each of the three MR head coils. 1a shows the tSNR maps corresponding to acquisition with a TMS coil placed in between the phantom and the MR head coils while 1b shows the tSNR maps for runs without a TMS coil placed on the phantom (TxRx and Magnetica). The tSNR scale is depicted at a range of 0-200 to emphasise differences between the three MR head coils. The maximum tSNR, mean and standard deviation is overlaid on each tSNR map. The TMS coil was placed on the right-hand side of the phantom with the approximate location marked next to each map. 1c shows subtraction maps for runs with a TMS coil subtracted from runs without a TMS coil (TxRx and Magnetica, brighter colours indicate greater tSNR loss).

To illustrate areas in the phantom with higher tSNR with the Magnetica and surface coils compared to the more standard TxRx coil, Figure 2 shows tSNR as a percentage of the TxRx coil. Of particular note are the areas close to the surface coils which range up to 600% tSNR of the TxRx, in line with the work from Navarro de Lara and colleagues (2015).

**Figure 2.**
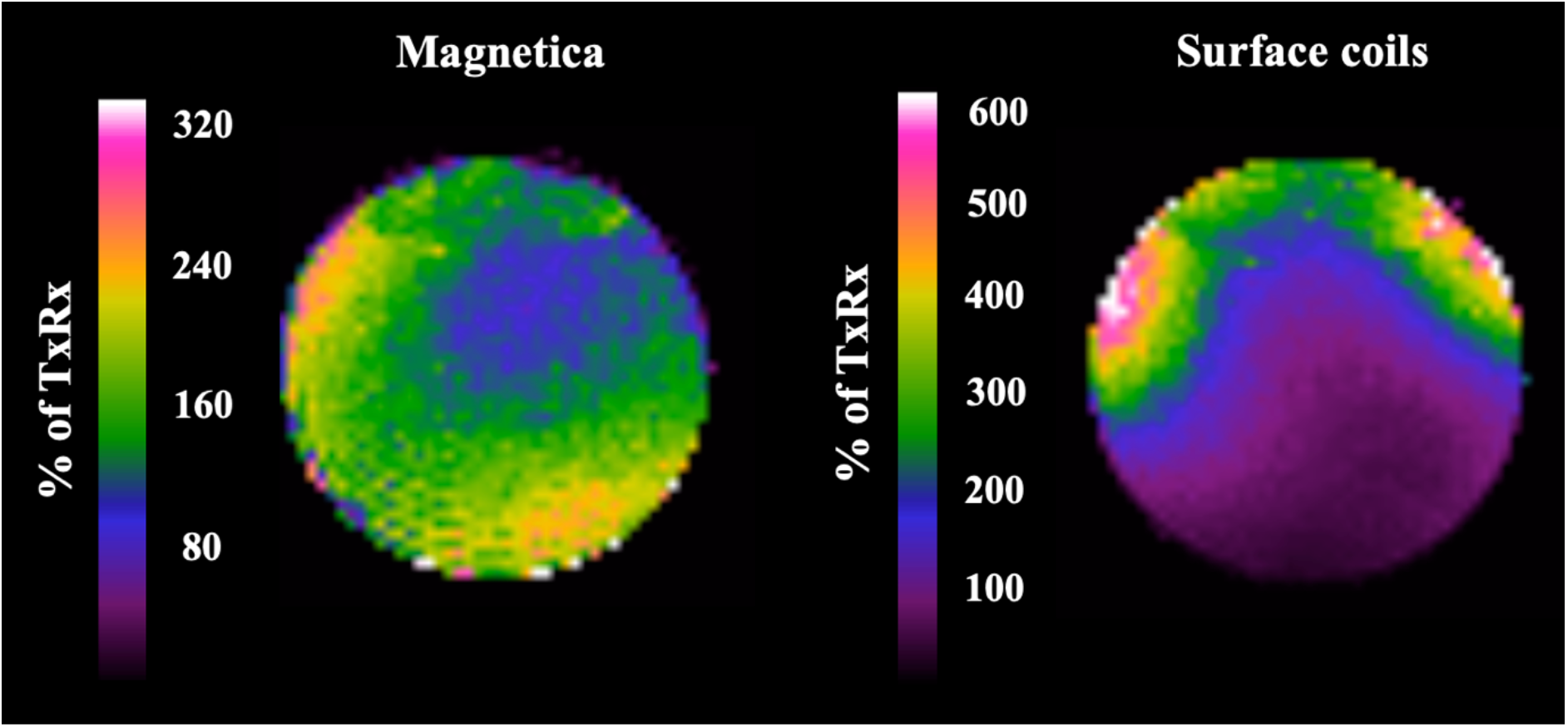
TSNR as a percentage of the TxRx coil, for both the Magnetica (left) and surface coils (right). Note the scale differs between the two MR coils.

## Discussion

These tSNR maps and corresponding data may be informative for researchers planning to conduct concurrent TMS-fMRI experiments. In previous work, the most common TMS-fMRI setup was with a standard bird cage coil, however, as shown here, better signal can be achieved with newer coils. The TMS-dedicated MR surface coils comparatively provide high resolution directly under the TMS coil (as demonstrated in previous work; de Lara et al., 2017) but lower resolution elsewhere not covered by the supplementary coil, reflecting considerable field inhomogeneity. Thus, there is a trade-off, and the choice of MR head coil will depend on whether the research aim is to focus on the result of stimulation at the targeted site and specific areas covered by the second surface coil, or whether the aim is to explore whole brain effects distant from the targeted region.

Asides from increased signal directly under the TMS coil compared to the other MR head coil setups, the dedicated surface coil solution can also be used with multiband imaging techniques and can potentially make in-bore neuronavigation more feasible. This solution also offers greater flexibility in terms of TMS coil placement, which is restricted with the other MR head coils within which space is limited. However, some coil arrangements may still not be possible because they would be uncomfortable for participants, and the position of the two surface coils may be challenging to standardise across participants. Another note is that higher stimulator output is needed with the surface coils, as the MR coil array is mounted below the TMS coil (Navarro de Lara et al., 2015; Bergmann et al., 2021). Thus, for participants with very high stimulation thresholds, suprathreshold intensities may not always be possible.

## Acknowledgements

This work was funded by Medical Research Council (UK) intramural funding SUAG/052/G101400. For the purpose of open access, the author has applied a Creative Commons Attribution (CC BY) licence to any Author Accepted Manuscript version arising from this submission.

## Data availability

Raw data is available on the Open Science Framework: https://osf.io/r6z2s/.

